# Inhibition of the neuromuscular acetylcholine receptor with atracurium activates FOXO/DAF-16-induced longevity

**DOI:** 10.1101/2020.07.24.219584

**Authors:** Rebecca L. McIntyre, Simone W. Denis, Rashmi Kamble, Michael Petr, Bauke V. Schomakers, Aldo Jongejan, Morten Scheibye-Knudsen, Riekelt H. Houtkooper, Georges E. Janssens

## Abstract

Transcriptome-based drug screening is emerging as a powerful tool to identify geroprotective compounds to intervene in age-related disease. We hypothesized that, by mimicking the transcriptional signature of the highly conserved longevity intervention of *FOXO3* (*daf-16* in worms) overexpression, we could identify and repurpose compounds with similar downstream effects to increase longevity. Our *in silico* screen, utilizing the LINCS transcriptome database of genetic and compound interventions, identified several FDA-approved compounds that activate FOXO downstream targets in mammalian cells. These included the neuromuscular blocker atracurium, which also robustly extends both lifespan and healthspan in *C. elegans*. This longevity is dependent on both *daf-16* signaling and inhibition of the neuromuscular acetylcholine receptor. Other neuromuscular blockers tubocurarine and pancuronium caused similar healthspan benefits. We demonstrate nuclear localization of DAF-16 upon atracurium treatment, and, using RNAseq transcriptomics, identify activation of DAF-16 downstream effectors. Together, these data demonstrate the capacity to mimic genetic lifespan interventions with drugs, and in doing so, reveal that the neuromuscular acetylcholine receptor regulates the highly conserved FOXO/DAF-16 longevity pathway.

## Introduction

Aging is a major risk factor for disease, and in recent years, cellular pathways and interventions influencing the aging process have steadily been uncovered. These are classified as the hallmarks of aging (López-Otín, Blasco, Partridge, Serrano, & Kroemer, 2013), which range from intracellular alterations, such as deregulated nutrient signaling pathways, to altered intercellular communication (López-Otín et al., 2013). Many genetic interventions have been identified to slow the progression of aging processes and finding pharmacological means to address this risk would be of great benefit to society (Figure 1A) (Flatt & Partridge, 2018; Fontana, Partridge, & Longo, 2010; Fulop, Larbi, Khalil, Cohen, & Witkowski, 2019; Longo et al., 2015; Partridge, Fuentealba, & Kennedy, 2020).

**Figure 1:**
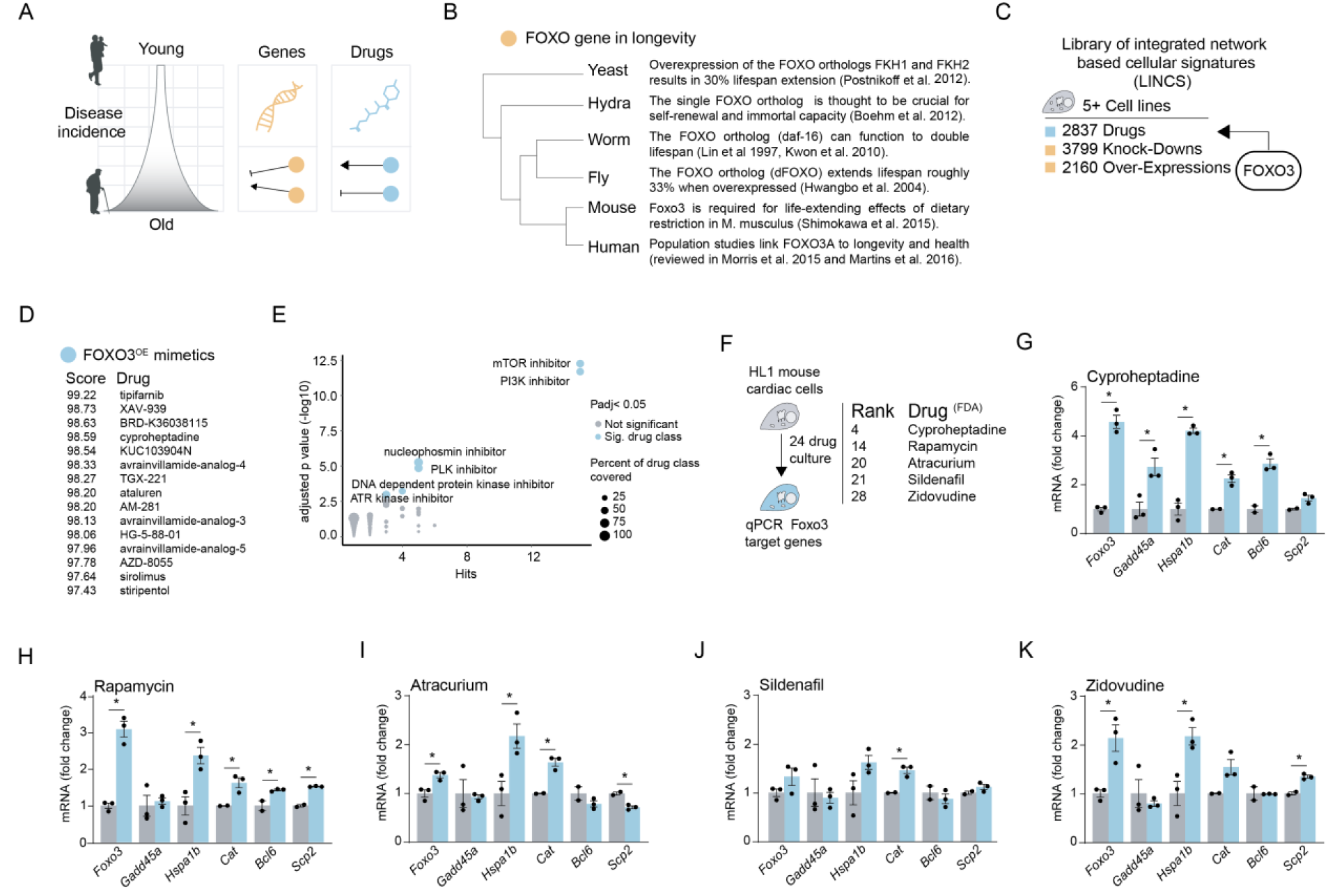
*In silico* drug screen identifies compounds that mimic transcriptional signature of *FOXO3* overexpression. A) Schematic depicting the increased incidence of disease throughout the aging process. Many genetic interventions (orange) have been described that intervene in the aging process to promote health and increase lifespan. Drug interventions (blue) are less prevalent, but here we describe efforts to identify compounds capable of producing the same effects. B) FOXO’s conserved role in longevity regulation across the phylogenetic tree. FOXO and its orthologs directly regulate, or are associated with, aging in organisms as distantly related as yeast and humans. C) The library of integrated network based cellular signatures (LINCS), a database that holds a core set of transcriptional signatures from at least five different commonly used cell lines. Transcriptional signatures for over 2500 drugs, over 3500 gene knock-downs and over 2000 gene over expressions are available. We used the *FOXO3* overexpression transcriptional signature present in LINCS to query for mimetic compounds. D) The top 15 hits mimicking *FOXO3* overexpression from the LINCS core compounds and their summary score integrating results from all cell lines tested. All compound hits can be found in Supplemental Table S1. E) Enrichment of drug classes in the candidate list, shows mTOR and PI3K inhibitors as highly enriched drug classes for their potential to mimic *FOXO3* overexpression among the candidate drugs. Names of drug classes are only displayed if passing significance of enrichment (adjusted *p* < 0.05, Y axis). Number of drug hits from the drug class is depicted on the X axis, while sizes of the dots in the scatterplot denote what percentage of the drug class were hits (i.e. 5 hits out of a class of 10 would denote a 50% coverage). F) Schematic of *in vitro* validation method to test candidate drugs. HL1 mouse cardiac cells were cultured with compounds for 24 hours of exposure. *FOXO3* and target gene expression was assessed by qPCR (mRNA level normalized to reference gene *Gapdh*). The top 5 ranked compounds from the candidate list (Figure 1D) that also had FDA approval status were used for testing. G-K) The expression levels of *Foxo3* and its target genes – *Gadd45a, Ccng2, Hspa1b, Cat, Bcl6, Scp2* as assessed by qPCR – in either control cells (grey) or cells treated with drug candidates (blue) – cyproheptadine (G), rapamycin (H), atracurium (I), sildenafil (J), and zidovudine (K). All drugs were assessed at a concentration of 50µM, except for rapamycin that was assessed at 10µM to avoid toxicity. Asterisk indicates significantly increased expression for each gene (p< 0.05, t-test) with the drug relative to control.

The hallmark of deregulated nutrient signaling includes many well-described longevity pathways, such as mechanistic target of rapamycin (mTOR) and Phosphatidylinositol 3-kinase (PI3K) (Houtkooper, Williams, & Auwerx, 2010). Integrated with both of these is the highly conserved genetic longevity intervention: activation or overexpression of the Forkhead Box O (FOXO) family of transcription factors. Overexpression of FOXO and its orthologs have been shown in yeast, worms, and flies to increase lifespan (Figure 1B) (Hwangbo et. al., 2004; Kwon et. al., 2010; Lin et. al., 1997; Postnikoff et. al., 2012). In hydra, FOXO is thought to be crucial for immortality (Boehm et al., 2012), and in mice it is required for the lifespan extending effects of dietary restriction (Figure 1B) (Shimokawa et al., 2015). In humans, the *FOXO3a* variant is specifically associated with longevity (Figure 1B) (Martins, Lithgow, & Link, 2016; Morris et. al., 2015).

Another hallmark of aging, altered intercellular communication, is most often discussed in relation to inflammation. Yet, more recently, the role of other forms of intercellular communication in the aging process have begun to emerge. One that is particularly relevant to this study is acetylcholine (ACh) signaling at the neuromuscular junction (NMJ). In *C. elegans*, ACh supplementation has been shown to improve stress tolerance (Furuhashi & Sakamoto, 2016). However, increased cholinergic transmission was shown to accelerate neurodegenerative phenotypes in mice (Sugita et al., 2016). There is currently very limited insight into the specific role of NMJ ACh signaling in aging and longevity, yet a connection may be present.

The identification of geroprotectors, compounds that can slow the aging process to delay the onset of age-related diseases, is a relatively new field. Despite the development of some high-throughput methods, screening for these compounds is time consuming, due to the need to evaluate full lifespans of model organisms (Moskalev et al., 2016). Though commonly used aging models such as the nematode *Caenorhabditis elegans* have relatively short lifespans compared to mammals, testing compounds on a large scale is still difficult.

An alternative to large scale *in vivo* screening is transcriptome-based *in silico* drug screening. Here, the majority of the screening process is performed computationally, thereby considerably narrowing down the candidate list of compounds requiring validation. This method, when applied to the transcriptional signature of the well-known longevity intervention, dietary restriction (DR), identified the DR mimetic allantoin (Calvert et al., 2016). Similarly, members of our group applied machine learning algorithms trained on human aging transcriptome datasets to drug-response transcriptomes. This method allowed the identification of HSP90 inhibitors, including monorden and tanespimycin, as geroprotectors (Janssens et al., 2019).

Here, using a transcriptomics-based *in silico* drug screen focused on the conserved genetic longevity intervention of overexpression of *FOXO3*, we identified a number of FDA-approved geroprotectors. From there, we focused on the neuromuscular blocker, atracurium, which activated FOXO targets in mammalian cell culture and extended both lifespan and healthspan in worms. This extension was dependent on *FOXO* (*daf-16* in worms) and antagonism of the acetylcholine receptor (AChR). Collectively, these results demonstrate a conserved longevity cascade in which inhibition of the AChR by atracurium leads to activation of FOXO/DAF-16 and therefore extended longevity.

## Results

### *In silico* drug screen identifies compounds that mimic transcriptional signature of *FOXO3* overexpression

In order to identify compounds that produce similar effects as *FOXO3* overexpression, we utilized the library of integrated network-based cellular signatures (LINCS), a database and software suite containing transcriptional signatures of both drug-treated human cell lines and genetic perturbations (Keenan et al., 2018; Subramanian et al., 2017). LINCS includes the transcriptional signatures of 2837 compounds, many of which are FDA-approved, originating from a core set of eight cell lines (PC3, VCAP, A375, HA1E, HCC515, HT29, MCF7, and HEPG2).

With the *FOXO3* overexpression transcriptional signature available in this database, we used the online software available within LINCS to search for compounds producing the most similar transcriptional effects to *FOXO3* overexpression (Figure 1C). For each compound, a standardized score was generated which ranged from -100 (highly opposing transcriptional signature to query candidate) to 100 (highly similar transcriptional signature to query candidate), and a summary score consolidating cell line data was generated for each compound. Summary scores greater than 90 were used as cutoff criteria to designate compounds with similar transcriptional signatures, as recommended by LINCS, and the ranked results were downloaded for further evaluation (Supplemental Table S1).

This approach generated a list of 129 candidate drugs mimicking *FOXO3* overexpression transcriptionally (Figure 1D; Supplemental Table S1). To categorize commonalities within this list, which contained both FDA approved drugs and compounds developed for research use, we used the drug descriptions associated with each compound to perform enrichment analysis. This allowed us to see which drug classes were most prevalent in our list, relative to the entire dataset. We found that both mTOR inhibitors and PI3Kinase inhibitors were strongly enriched as drug classes that matched the *FOXO3* overexpression mRNA signature (Figure 1E). Because mTOR and PI3Kinase inhibitors are both known to increase lifespan, and act within the FOXO longevity pathway (Babar et. al., 1999; Hay, 2011; Johnson, Rabinovitch, & Kaeberlein, 2013; Moskalev & Shaposhnikov, 2010), we concluded that our approach could indeed identify compounds that are known to extend lifespan through the FOXO signaling pathway.

As it is more straightforward to repurpose drugs already used in a clinical setting than to develop new ones, we continued investigation with our top five hits from the screen that already possessed FDA approval (Figure 1F). These compounds included two positive controls known to extend model organism lifespan: the serotonin antagonist cyproheptadine (ranked 1st on the FDA-approved subset; ranked 4th overall) (Petrascheck, Ye, & Buck, 2007) and the mTOR inhibitor rapamycin (2nd FDA-approved; 14th overall) (Robida-Stubbs et al., 2012; Xu et al., 2018) and three other compounds; the neuromuscular blocker atracurium (3rd FDA-approved; 20th overall), the vasodilator sildenafil (4th FDA-approved; 21st overall), and the nucleoside reverse transcriptase inhibitor (NRTI) zidovudine (5th FDA-approved; 28th overall).

We next sought to investigate these individual compounds’ ability to activate the FOXO pathway *in vitro*. Because FOXO has long been implicated in cardiac aging (Wong & Woodcock, 2009), we cultured mouse HL1 cardiac cells with each compound individually for 24 hours and assessed the gene expression of *Foxo3* itself and known *Foxo3* target genes (Figure 1F). Target genes included *Gadd45a*, a DNA repair gene directly stimulated by FOXO3 activity (Tran et al., 2002), *Ccng2* and *Bcl6*, both cell cycle related genes also directly regulated by FOXO3 (Fernandez de Mattos et al., 2004; Fu & Peng, 2011), the catalase *Cat* and the sterol carrier *Scp2*, which are part of the cellular antioxidant defense system regulated by *Foxo* transcription factor activity in general (Dansen et al., 2004; Klotz et al., 2015), and *Hspa1a*, which encodes a heat shock protein regulated by FOXO transcription factors, conserved across species (Webb, Kundaje, & Brunet, 2016). Relative to untreated cells, we found that most compounds could significantly activate the expression of *Foxo3* itself as well as various *Foxo3* target genes (Figure 2G-K). These data suggest that our compound screen was successful in identifying compounds that, at least transcriptionally, mimic an induction of *FOXO3*.

**Figure 2:**
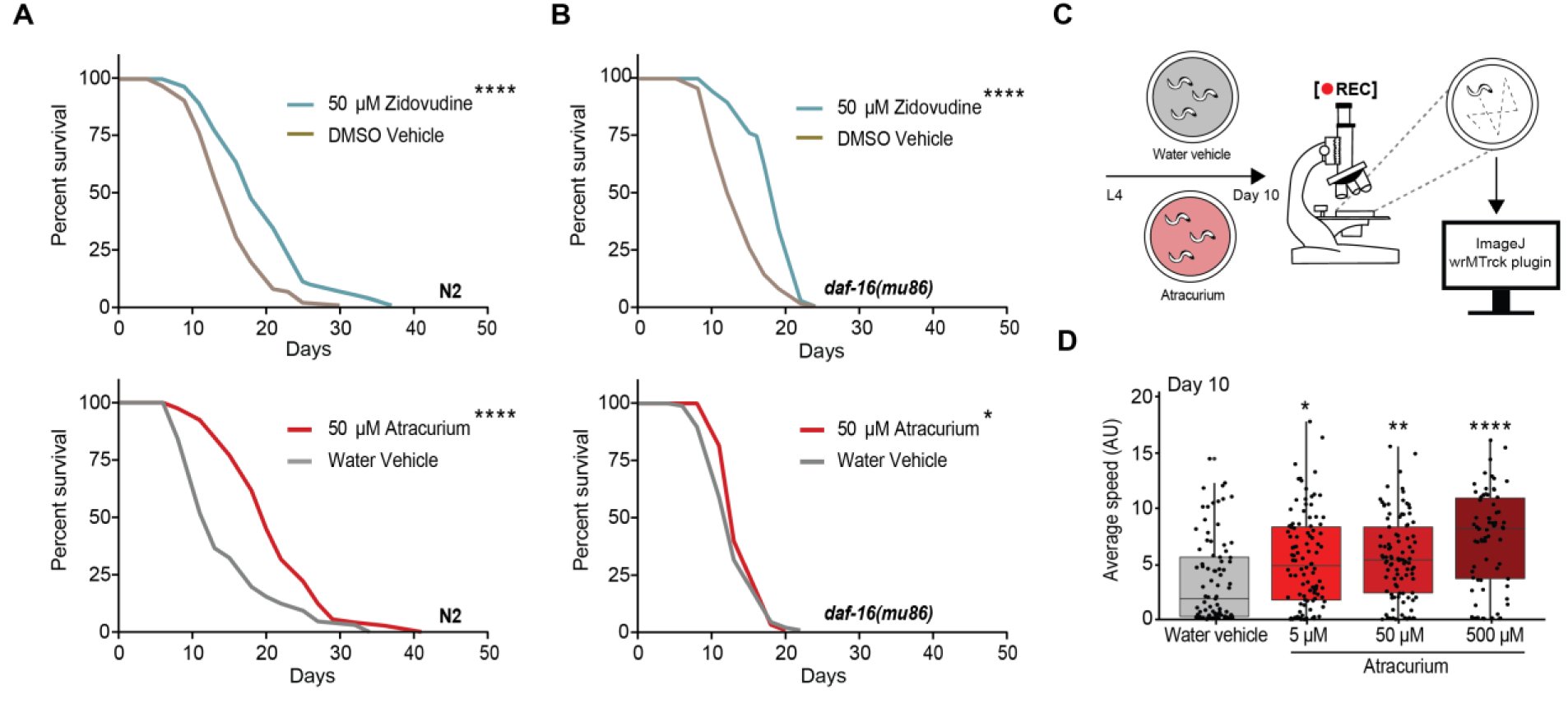
Zidovudine and atracurium extend lifespan in *C. elegans*, but only atracurium depends on *daf-16*. A) Survival curves showing that zidovudine and atracurium (both 50 μM) extend lifespan in wild type (N2) *C. elegans*. B) Survival curves showing that zidovudine, but not atracurium (both 50 μM), extends lifespan in the *daf-16(mu86)* strain. All statistical comparisons of survival curves are determined by log-rank tests. For each condition, n=100 worms. See Supplemental Table S2 for lifespan statistics. ****p<0.0001, *p<0.05. C) Schematic describing the workflow for testing healthspan using age-related mobility in *C. elegans*. Worms are cultured on NGM supplemented with either water vehicle or atracurium beginning at the L4 stage. At day 10 of adulthood, ∼50 worms per condition are moved to plates without bacteria and are filmed through a microscope after stimulation. These videos are processed with ImageJ and average crawling speed is measured by the wrMTrck ImageJ plugin. D) Mobility of atracurium-treated N2 worms at day 10 of adulthood. Atracurium improves healthspan in a dose-dependent manner (5 μM, 50 μM, 500 μM). For each condition, n=∼70-100 measurements of ∼50 worms. Statistical significance is determined by a one-way ANOVA, and represented p-values are each compared to the water vehicle. ****p<0.0001, **p<0.01, *p<0.05.

### Zidovudine and atracurium both extend lifespan in *C. elegans*, but only atracurium depends on *daf-16*

To test the effects of the identified compounds on lifespan, we turned to *C. elegans*, a simple and well-described model organism for aging. As noted above, cyproheptadine and rapamycin are known lifespan extenders (Petrascheck et al., 2007; Robida-Stubbs et al., 2012; Xu et al., 2018), so we proceeded to test lifespan effects of atracurium, sildenafil and zidovudine. Of these three, zidovudine and atracurium robustly and reproducibly extended lifespan at a concentration of 50 μM (Figure 2A; Supplemental Table S2), whereas sildenafil led to no consistent difference in lifespan with the concentration and conditions we tested (Supplemental Table S2).

Because our compound screen was based on overexpression of *FOXO3*, we hypothesized that lifespan extension from our positive hits would be dependent on *daf-16*. We therefore tested whether zidovudine and atracurium extend lifespan in the *daf-16(mu86)* mutant strain. Zidovudine maintained a robust lifespan extension in the mutant, while the lifespan extension induced by atracurium was almost completely abrogated (Figure 2B). Together these data suggest that, despite similar transcriptional profiles, atracurium is dependent on *daf-16*, while zidovudine is not. We expect that the highly conserved nature of *FOXO/daf-16* may be of significant translational relevance and therefore continued investigation of atracurium to further understand the upstream signaling at play.

### Atracurium extends healthspan in *C. elegans*, as measured by age-related mobility

Healthy aging is not only determined by lifespan, but also healthspan. One way to measure the health of worms is through mobility, or average crawling speed (Pierce-Shimomura et al., 2008). In order to assess the healthspan of atracurium-treated worms, we tested their mobility at day 10 of adulthood, as this is generally the point at which the majority of control worms are still alive, but experience mobility impairment. We recorded the movement of a sample of worms, and used an image processing platform to analyze the films (Figure 2C) (Nussbaum-Krammer et.al., 2015). We titrated the concentration of atracurium to additionally test healthspan for dose-response. At day 10, atracurium significantly increased age-related mobility in a dose-dependent manner (Figure 2D). These data suggest that atracurium extends not only lifespan, but healthspan as well.

### Atracurium extends healthspan in *C. elegans* by antagonizing the neuromuscular acetylcholine receptor

Atracurium is a neuromuscular blocker and is used in patient care as an anesthetic muscle relaxant. We therefore considered the possibility that, if the pharyngeal muscles were relaxed by atracurium treatment, the worm would be unable to pump bacteria efficiently into its body, consequently reducing food intake and inducing a state of dietary restriction (DR). We therefore tested pharyngeal pumping on day 1 of adulthood. Contrary to our expectation, worms treated with atracurium showed a significantly increased rate of pumping when compared to their control counterparts (Figure 3A). Also contrary to a DR phenotype, the body area of the worms at day 10 of adulthood was increased by atracurium relative to untreated worms in a dose-dependent manner (Figure 3B). These data together suggest that DR is not the cause of atracurium’s extension of healthspan.

**Figure 3:**
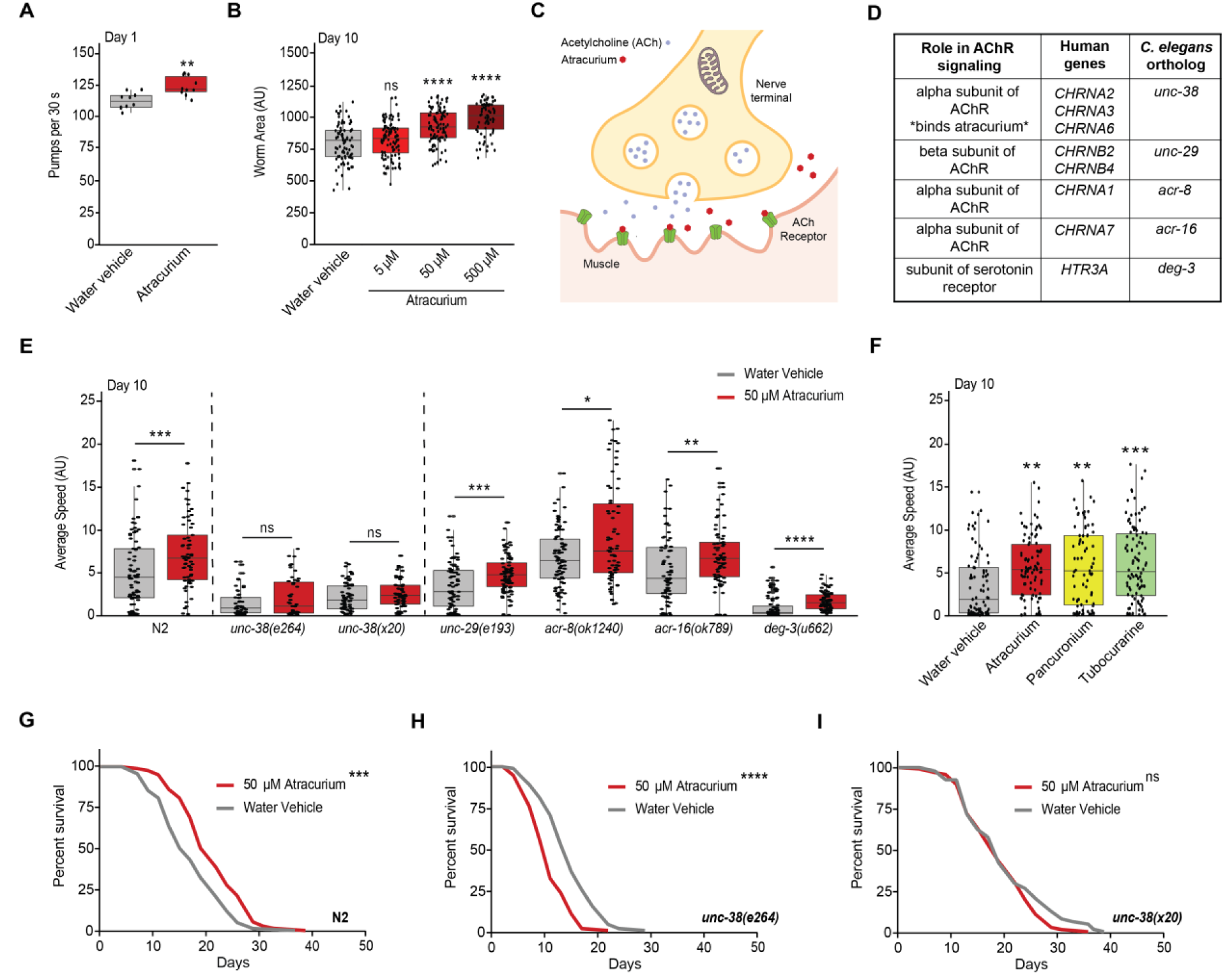
Atracurium extends healthspan by antagonizing the neuromuscular acetylcholine receptor. A) Pharyngeal pumping at day 1 of adulthood in control and atracurium (50 μM) treated N2 worms. Despite its clinical function as an anesthetic muscle relaxant, atracurium does not reduce pumping. For each condition, n=10 worms. Statistical significance is determined by unpaired t-test. **p<0.01 B) Body size of atracurium-treated N2 worms at day 10 of adulthood. Increased concentration of atracurium causes increased body size in a dose-dependent manner (5 μM, 50 μM, 500 μM). For each condition, n=∼70-100 measurements of ∼50 worms. Statistical significance is determined by a one-way ANOVA, and represented p-values are each compared to the water vehicle. ****p<0.0001, ns - not significant C) Representation of the canonical mechanism of atracurium at the neuromuscular junction (NMJ) relevant to this study. In humans, atracurium is used to relax muscles, and does so by antagonizing the acetylcholine receptor (AChR). Under normal conditions, acetylcholine (ACh) binds the receptor, causing depolarization of the channel, and contraction of the muscle. Atracurium binds the AChR in place of ACh, preventing this depolarization and contraction. D) Table describing orthogolous ACh signaling genes in humans and *C. elegans*. In humans, atracurium binds the AChR alpha subunit encoded by the *CHRNA2* gene. E) Mobility of atracurium-treated (50 μM) N2 worms and mutants of players in NMJ ACh signaling at day 10 of adulthood. Atracurium improves healthspan in N2 worms, but this effect is abolished in worms with mutant *unc-38*. The other ACh pathway mutants are still responsive to atracurium, suggesting that the increased healthspan is dependent on this specific subunit of the AChR. For each condition, n=∼50-90 measurements of ∼50 worms. Statistical significance for each mutant is determined by an individual Mann-Whitney U test comparing treated and untreated. ****p<0.0001, ***p<0.001, **p<0.01, *p<0.05, ns - not significant. F) Mobility at day 10 of adulthood of N2 worms treated with atracurium, tubocurarine or pancuronium (all 50 μM). All compounds improve healthspan. For each condition, n=∼70-100 measurements of ∼50 worms. Statistical significance is determined by a one-way ANOVA, and represented p-values are each compared to the water vehicle. ***p<0.001, **p<0.01. G) Control survival curves (performed in parallel to those in panels H and I) showing that atracurium (50 μM) extends lifespan in wild type (N2) *C. elegans*. ***p<0.001. H) Survival curves showing that atracurium (50 μM) does not extend lifespan in the *unc-38(e264)* mutant strain (atracurium shortens lifespan ****p<0.0001). I) Survival curves showing that atracurium (50 μM) does not extend lifespan in the *unc-38(x20)* mutant strain. For each condition, n=100. Lifespan statistics are determined by log-rank tests. See Supplemental Table S2 for lifespan statistics.

At the NMJ in physiological conditions, ACh crosses the synapse to bind to the AChR, causing the channel to depolarize and the muscle to contract (Rand, 2007). Atracurium competitively antagonizes the AChR (Lee, 2001; Wishart et al., 2018), preventing this contraction and allowing the muscle to relax instead (Figure 3C). We questioned if atracurium extends healthspan through this pathway, or instead through an off-target effect. Atracurium binds an alpha subunit of the AChR, encoded by *CHRNA2* in humans (Wishart et al., 2018) The closest homolog to this subunit in *C. elegans* is encoded by the gene *unc-38* (Figure 3D) (Fleming et al., 1997; Harris et al., 2020). There are multiple strains of *C. elegans* available with mutant *unc-38*, so we chose to utilize both *unc-38(e264)* and *unc-38(x20)*. Additionally, as the AchR system is not entirely homologous between human and *C. elegans* (Rand, 2007), we also included a range of mutants of genes involved NMJ signaling. There are 27 genes identified AChR genes in *C. elegans*, which have been divided into five classes of protein subunits based on sequence similarity (Rand, 2007). We wished to test all five classes, and therefore included mutants for *unc-29(e193), acr-8(ok1240), acr-16(ok789)*, and *deg-3(u662)* (Figure 3D). To determine if the effects we saw with atracurium were dependent on any of these subunits, we tested healthspan at day 10 of adulthood. We saw again increased healthspan in wild type N2 worms treated with atracurium compared to those untreated (Figure 3E). However, in both *unc-38(e264)* and *unc-38(x20)* mutant worms, atracurium was no longer able to prolong healthspan (Figure 3E). This effect was specific to *unc-38*, as atracurium was still able to increase healthspan in the other four NMJ mutants tested (Figure 3E). Therefore, these data show that *unc-38* is required for atracurium-induced improvements in healthspan.

Atracurium is one of many neuromuscular blocking compounds used in human anesthesia (Lee, 2001). We questioned if other drugs that acted on the same pathway would also lead to extension of healthspan in *C. elegans*. To investigate this, we exposed worms to two of these compounds that were readily commercially available: pancuronium and tubocurarine, both of which act on the same alpha subunit of the AChR (Lee, 2001; Wishart et al., 2018). At day 10 of adulthood, these compounds extended healthspan compared to untreated worms, similar to the effects of atracurium (Figure 3F). These data suggest that the healthspan effects we see with atracurium are not specific to atracurium alone, but instead can be generalized to neuromuscular blockers that antagonize the alpha subunit of NMJ AChR.

Having determined that healthspan of the *unc-38* mutant worms was unaffected by atracurium, and that blocking this subunit with more compounds created similar healthspan benefits, we tested if absence of *unc-38* also prevented atracurium-induced lifespan extension. Indeed, unlike treatment in N2 worms (Figure 2A, 3G), atracurium was unable to extend lifespan in either of these mutants (Figure 3H,I). In fact, in the *unc-38(e264)* mutant, atracurium appeared to have detrimental effects on lifespan (Figure 3H). Altogether, these data suggest that atracurium specifically requires the *unc-38* subunit of the AChR to enact improvements on healthspan and lifespan in *C. elegans*, and these effects are not due to an off-target response.

### Atracurium stimulates nuclear translocation of DAF-16

Having determined that the lifespan extension induced by atracurium is dependent on *daf-16*, we next aimed to validate that atracurium treatment causes physical translocation of *daf-16* to the nucleus. To investigate localization of DAF-16, we employed a transgenic *C. elegans* strain that expresses DAF-16 tagged with GFP (Zhao et al., 2017). Heat stress activates DAF-16, so we utilized it as a positive control for DAF-16 localization to nuclei of neurons within the heads of the mutant worms (Lin et al., 2018; Figure 4A,B). Similarly, atracurium treatment for 24 hours (from L4 stage to day 1 of adulthood) caused significantly increased nuclear localization of DAF-16 in worms when compared to those untreated (Figure 4A,B). Chi-square analysis generated a p-value of 0 for both heat stress and atracurium when each was compared to the water vehicle control condition (Figure 4B). Therefore, we conclude that atracurium’s inhibition of the UNC-38 subunit causes nuclear localization of DAF-16.

**Figure 4:**
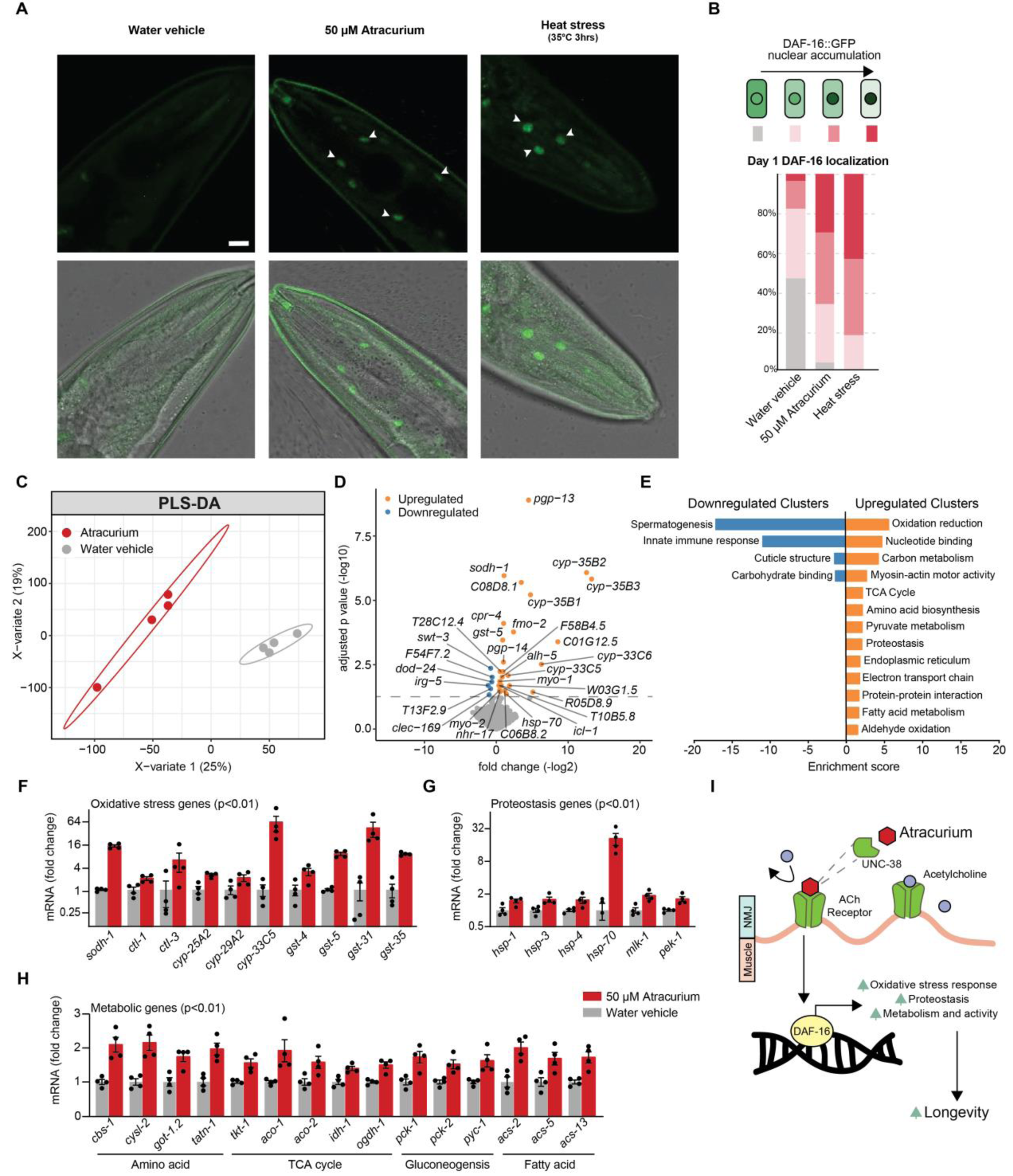
Atracurium stimulates translocation of DAF-16 to the nucleus and upregulation of DAF-16 downstream targets. A) Representative images of heads of worms expressing DAF-16::GFP upon treatment with water vehicle, atracurium (50 μM). Treatments are compared to heat stress at 35°C for 3 hours as a positive control. Images were taken on day 1 of adulthood. The upper panel shows fluorescent images of DAF-16::GFP nuclear localization, while the lower panels display overlay of DAF-16::GFP fluorescent images with bright field images of worms. Scale bar measures 10 μM and can be applied to all images. White arrows indicate neuronal nuclei. B) Quantification of DAF-16 nuclear localization in A. n=∼20-30 different animals, pooled from two independent experiments. Atracurium treatment significantly increases the proportion of worms with nuclear-localized DAF-16, similar to the heat stress positive control. A chi-square test of independence was performed to assess the significance of differences in the levels of DAF-16 nuclear enrichment between each of the treatments compared to water vehicle control (p = 0 for both comparisons). C) Partial least squares-discriminant analysis (PLS-DA) showing group separation based on differentially expressed genes in worms treated with atracurium (50 μM) compared to water vehicle control worms. D) Differential analysis showing significantly up- and downregulated genes (using adjusted p-value<0.05). RNAseq results can be found in Supplemental Table S3. E) Functional annotation clustering of the significantly (using unadjusted p-value<0.01) up- and downregulated genes performed using the DAVID Bioinformatics Database with an enrichment score >1.3. F-H) Known DAF-16-regulated oxidative stress response (F), proteostasis (G), and metabolic genes (H) that are upregulated with atracurium treatment, as indicated by RNAseq. For each fold change shown, unadjusted p-value is <0.01.I) Hypothesized mechanism for atracurium-mediated longevity. We have shown that atracurium requires the AChR subunit UNC-38 to extend healthspan and lifespan, thereby competitively antagonizing the receptor and this inhibition causes DAF-16-mediated lifespan extension. Longevity is likely caused by DAF-16 activation of genes known to induce longevity such as those involved in protective oxidative stress and proteostasis responses, as well as metabolic remodeling.

### RNA sequencing demonstrates induction of protective stress responses and metabolic remodeling by FOXO/DAF-16 downstream effectors

Having concluded that atracurium causes nuclear localization of DAF-16, we wished to examine the downstream transcriptional effects. Therefore, we performed RNA sequencing on worms treated for 24 hours (from L4 stage to day 1 of adulthood) with atracurium compared to those untreated. The two treatment groups were readily distinguishable and separable using partial least squares discriminant analysis (PLS-DA; Figure 4C). Differential analysis showed a number of significant hits after correcting for multiple hypothesis testing (Figure 4D). We noted that many of the significantly upregulated genes are FOXO downstream targets, for instance the alcohol dehydrogenase *sodh-1*, the heat shock protein *hsp-70*, and several members of the cytochrome P450 family (Figure 4D).

In order to form a more comprehensive view of the changing processes, we investigated the differentially expressed genes further, using the Database for Annotation, Visualization and Integrated Discovery (DAVID) bioinformatics resource (Huang, Sherman, & Lempicki, 2009). We performed functional annotation clustering, using a less strict significance cutoff (unadjusted p<0.01). Enriched annotation clusters included functional categories, GO-terms, KEGG pathway features and protein domains (Supplemental Table S4). Upregulated clusters included protective stress responses, for instance those relating to oxidative stress and proteostasis, as well as a number of metabolic processes, such as carbon metabolism and the TCA cycle (Figure 4E).

Activation of oxidative stress responses and proteostasis, as well as modulation of metabolic processes, are also key downstream effects of DAF-16 activation, and the majority of genes within the clusters we identified are also known to be upregulated by DAF-16 (Harris et al., 2020; Honda & Honda, 1999; Li & Zhang, 2016; McElwee, Bubb, & Thomas, 2003; Webb et al., 2016; Zečić & Braeckman, 2020). Significantly upregulated oxidative stress response genes included *sodh-1*, catalase genes *ctl-1* and *ctl-3*, and members of the cytochrome P450 and glutathione S-transferase families (Figure 4F), all of which are DAF-16 targets (Webb et al., 2016). Similarly, a number of DAF-16-dependent heat shock proteins were upregulated, as well as additional genes involved in protein scaffolding and unfolded protein response (Figure 4G). Finally, various aspects of cellular metabolism were upregulated upon atracurium treatment, including DAF*-*16-dependent genes involved in amino acid metabolism, the TCA cycle, gluconeogenesis, and fatty acid metabolism (Figure 4H). Together these data confirm that the longevity induced by atracurium’s competitive antagonism of the AChR is indeed mediated by DAF-16 activation, likely through metabolic remodeling and the induction of protective stress responses (Figure 4I).

### Discussion

Increasing evidence demonstrates that slowing the progression of aging processes with geroprotective compounds can treat age-related disease, thereby necessitating the discovery and deeper understanding of geroprotectors (Moskalev et al., 2016; Partridge et al., 2020). We have utilized an *in silico* drug screening strategy to identify compounds that mimic the *FOXO3* overexpression transcriptome. In doing so, we identified a number of FDA-approved compounds, both known and unknown previously to defer aging. The top five compounds of the screen activated *Foxo3* targets in mammalian cell culture, at least in part. When we tested for longevity effects *in vivo*, both the NRTI zidovudine and the neuromuscular blocker atracurium robustly extended lifespan in *C. elegans*, but only atracurium was dependent on *FOXO*/*daf-16*. Atracurium extended not only lifespan, but also healthspan in worms. We found that these healthspan effects were dependent on the AChR subunit encoded by *unc-38*, the closest worm homolog to human *CHRNA2*, which encodes the subunit known to bind atracurium upon treatment. Microscopy demonstrated DAF-16 nuclear localization, and RNA sequencing revealed differential expression of genes known both to be regulated by DAF-16 signaling and to modulate aging.

We chose to base our *in silico* screen on overexpression of *FOXO3*, as it is one of the most widely recognized and conserved genetic interventions to promote longevity (Martins et al., 2016). We therefore hypothesized that compounds mimicking this transcriptome, particularly by activating FOXO/DAF-16 itself, would also induce conserved longevity effects. Specifically, *FOXO3* expression has been associated with long-lived populations of humans in several independent cohorts (Broer et al., 2015; Morris et. al., 2015; Soerensen et al., 2015) suggesting that activation of this pathway is one of the most promising routes to benefiting human age-related diseases.

One of our hits *in vivo*, however, did not follow this expectation: zidovudine robustly extended lifespan, but was not dependent on *daf-16*, suggesting that its effects on gene expression— inducing a FOXO-like transcriptome—are caused by factors acting downstream of the transcription factor. Effects of NRTIs have been studied previously in *C. elegans* and are hypothesized to inhibit mitochondrial DNA (mtDNA) polymerase γ function, leading to a depletion of mtDNA, and decreased mitochondrial respiration due to a lack of subunits of the respiratory chain (Brinkman, ter Hofstede et. al., 1998; Lewis & Dalakas, 1995). However, *C. elegans* exposed to zidovudine have relatively modest reductions in mtDNA and yet simultaneously experience diminished oxygen consumption and alterations to mitochondrial morphology (de Boer et al., 2015). Impairment of mitochondrial function is a well-established lifespan extension mechanism (Cho, Hur, & Walker, 2011; Dillin et al., 2002; Houtkooper et al., 2013; Lee et al., 2003; Liu et al., 2020; Molenaars et al., 2020), so we expect that this is also the mechanism by which zidovudine extends lifespan in worms. While this mechanism is not the focus of our current study, we remain intrigued by the fact that FOXO’s longevity signature is potentially activated through other means. It would be interesting in the future to use zidovudine as a model for the translation potential of activating other FOXO-like longevity pathways.

The healthspan and lifespan extensions induced by atracurium were dependent on the AChR subunit *unc*-38, demonstrating that atracurium works to promote longevity through its canonical mechanism. There is sparse previous work investigating the role of acetylcholine signaling in aging. However, in one recent study, mice with overexpression of the vesicular acetylcholine transporter (VAChT) allowed for analysis of how increased release of ACh affected development and aging (Sugita et al., 2016). In this model, increased ACh accelerated aging of the NMJ, measured by fragmentation, denervation, sprouting of motor axon nerve endings, and innervation by multiple motor axons (Sugita et al., 2016). Degeneration of the NMJ preceded motor deficits and muscular atrophy in the overexpression mice, and, upon crossing mice of this genotype with a mouse model for ALS, increased ACh transport led to accelerated pathology and early death (Sugita et al., 2016). These data provide insight into the converse mechanism—over activation of the AChR as a mechanism for premature aging—and align with our findings that inhibition of the AChR promotes longevity.

The lifespan extension caused by atracurium was also dependent on *daf-16*, and the drug indeed caused increased DAF-16 nuclear localization. Our RNAseq data reinforced this finding, as a significant number of differentially expressed genes in atracurium-treated worms compared to controls were known downstream effectors of DAF-16. Altogether, these data suggest that inhibition of acetylcholine signaling at the neuromuscular junction causes activation of DAF-16 and the transcription of genes within its lifespan-extending pathways. However, how DAF-16 is activated downstream of ACh signaling is still unclear. Identifying these intermediate steps would be an interesting focus of future study. Particularly when we imagine developing pharmacological agents that could be used in a clinical setting, finding targets directly upstream of FOXO would be distinctly appealing.

Altogether, our results demonstrate the power of *in silico* drug screening, particularly the potential to mimic genetic longevity interventions with compounds. In doing so, we identified the neuromuscular blocker atracurium as a geroprotector, a compound that activates FOXO3 targets *in vitro* in mammalian cell culture, and promotes longevity *in vivo* in worms. Using atracurium, we show that neuromuscular acetylcholine signaling acts as a regulator of the conserved FOXO longevity pathway. This finding facilitates deeper understanding of this aging mechanism and subsequent identification of potential targets for the treatment of age-related diseases.

## Experimental procedures

### LINCs database compound screen

The online library of integrated network-based cellular signatures (LINCS) (Keenan et al., 2018; Subramanian et al., 2017) was accessed (September 2017) through the cloud-based software platform CLUE (https://clue.io/). The core dataset termed ‘touchstone,’ containing transcriptional signatures from eight different cell lines (PC3, VCAP, A375, HA1E, HCC515, HT29, MCF7, HEPG2) of 2837 different drug treatments, 3799 different gene knock-downs, and 2160 different gene over-expressions was used. From these, the FOXO3^oe^ transcriptional signature (Broad ID: ccsbBroad304_00577) was used as a query base to search for compounds with similar transcriptional signatures, and ranked results including a summary score consolidating cell line data, ranging from -100 (opposing signature) to 100 (mimicking signature) were downloaded as.gct files (version 1.3). These ranked results for all cells and their summary score are available in Supplemental Table S1. A cutoff was applied to the ranked list whereby compounds with a score greater than 90 were considered to match the FOXO3^oe^ transcriptional signature. The top 5 FDA approved drugs within this list were used for further *in vitro* and *in vivo* evaluation.

### Cell culture

HL1 cardiomyocytes were cultured in 0.02% gelatin (Sigma-Aldrich; Darmstadt, Germany) coated flasks in Claycomb Medium (Sigma-Aldrich) supplemented with 2 mM L-glutamine (Gibco; Dublin, Ireland), antibiotic mixture of 100 U/mL penicillin and 100 µg/mL streptomyocin (Lifetechnology; Bleiswijk, The Netherlands), 0.25 µg/mL fungizone (Lifetechnology), and 10% (v/v) fetal bovine serum (FBS) (Bodinco; Alkmaar, The Netherlands) at 37°C in a humidified atmosphere of 5% CO_2_.

Cells were cultured for 24 hours with the compounds described -50 µM cyproheptadine hydrochloride sesquihydrate (Sigma Aldrich), 10 µM rapamycin (Selleckchem; Munich, Germany), 50 µM atracurium besylate (Sigma Aldrich), 50 µM sildenafil citrate salt (Sigma Aldrich), 50 µM zidovudine (Sigma Aldrich) - for 24 hours before isolation of RNA.

### Extraction of mRNA from cells and quantitative real-time PCR (qPCR)

Isolation of total mRNA from cells was performed with TRI-reagent (Sigma-Aldrich), and 1 µg of extracted RNA was reverse transcribed into cDNA according to manufacturer’s instructions using the QuantiTect Reverse Transcription Kit (QIAGEN; Venlo, The Netherlands). Quantitative gene expression analysis was performed using the LightCycler® 480 SYBR Green I Master (Roche; Woerden, The Netherlands) and measured using the LightCycler® 480 Instrument II (Roche). Gene-specific primers were synthesized according to the sequences in Supplemental Table S5. The N0 values of target genes were normalized to the reference gene *Gapdh*. All experiments were performed in triplicate. Statistical analysis compared fold change in gene expression relative to the mean value of controls between treated and untreated HL1 cells using a t-test in Graphpad Prism.

### Worm strains and maintenance

*C. elegans* strains N2 Bristol, *daf-16(mu86), unc-38(e264), unc-38(x20), unc-29(e193), acr-8(ok1240), acr-16(ok789), deg-3(u662), daf-16(mu86);muIs61* and *E. coli* strain OP50 were obtained from *Caenorhabditis* Genetics Center (CGC; University of Minnesota, Minneapolis, MN, USA). Worms were routinely grown and maintained on nematode growth media (NGM) agar plates seeded with OP50 *E. coli* at 20 °C as previously described (Brenner, 1974).

### Pharmacological treatment of *C. elegans*

All chemicals were obtained from Sigma Aldrich. Atracurium besylate, laudanosine, tubocurarine hydrochloride pentahydrate and pancuronium bromide were dissolved in water at a concentration of 16.67 mM. Zidovudine and sildenafil citrate salt were dissolved in DMSO at a concentration of 50 mM. These compounds were added to plates just before pouring at the concentrations described. For all experiments, worms were treated with compounds from L4 stage onwards, and, unless otherwise described, plates were changed once a week to ensure consistent exposure to the compound.

### Lifespan analysis

Gravid adult worms were age-synchronized using alkaline hypochlorite treatment, and incubated in M9 buffer overnight. L1 larval stage worms were seeded to NGM plates and grown to L4 stage. At L4 stage, worms were transferred to plates supplemented with compounds as described and 10 µM 5-fluorouracil (Sigma Aldrich). 10 µM 5-fluorouracil treatment continued for the first two weeks of life. All assays were performed at 20°C, and the L4 stage was counted as day 0 of life. One hundred worms were used per condition in every lifespan experiment. Survival was analyzed every other day and worms were considered dead when they did not respond to repeated prodding. Worms that were missing, displaying internal egg hatching, losing vulva integrity, and burrowing into NGM agar were censored. Statistical analyses of lifespan were calculated by Log-rank (Mantel-Cox) tests on Kaplan-Meier curves in GraphPad Prism.

### Healthspan measurements

Gravid adult worms were age-synchronized using alkaline hypochlorite treatment, and incubated in M9 buffer overnight. L1 stage worms were seeded to NGM plates. Worms were transferred to plates supplemented with compounds and 10 µM 5-fluorouracil (Sigma Aldrich) at the L4 larval stage. All assays were performed at 20°C, and the L4 stage was counted as day 0 of life. Plates were changed twice per week to maintain exposure to the compounds. All functional assays were performed at least twice, one of which is represented in the data shown.

#### Mobility analysis and body size

At day 10 of adulthood, ∼50 worms were transferred to NGM plates without OP50, stimulated by tapping the plate, and immediately recorded for 200 cycles at room temperature using a Leica (Amsterdam, The Netherlands) M205 FA fluorescent microscope and Leica DFC 365 FX camera. Images were captured using Leica Application Suite X software, then processed with the wrMTrck plugin for ImageJ (Nussbaum-Krammer et al., 2015). Measurements of body size were also derived from these analyses. Data from wrMTrck were analyzed and visualized using a custom script in R v3.6.3 (R Core Team, 2013). Statistical analysis compared conditions to their respective control with a one-way ANOVA corrected for multiple testing.

#### Pharyngeal pumping

At day 1 of adulthood, worms were visualized at room temperature using a Leica M205 FA fluorescent microscope. Pumps were counted by eye for 30 seconds, and 10-15 worms per condition were tested. Statistical analysis was performed in R v3.6.3 (R Core Team, 2013), comparing the two conditions with t-testing.

### Microscopy

Mutant worms expressing DAF-16::GFP were synchronized, grown and treated with compounds from L4 stage as described. To test DAF-16 nuclear localization, day 1 animals were immobilized for 30 seconds in 40 mM levamisole (Santa Cruz Biotech) in M9 buffer and mounted on 2% agarose pads. Nuclear localization of DAF-16::GFP was visualized using a Leica DMI6000 inverted confocal microscope containing a 40x, 1.30 Oil CS2 objective lens and a Leica TCS SP8 SMD camera. Images were captured using Leica Application Suite X software. All samples were imaged at room temperature. To avoid subtle localization caused by starvation, mounting and imaging conditions, all photomicrographs were taken within 5 minutes of mounting. To prevent interference of worm intestinal autofluorescence, images were focused on neuronal nuclei in the head. For heat stress, worms were exposed to 35°C for 3 hours. Images were taken of 20-30 worms per condition over two independent experiments and results are compiled from both replicates. Quantification of nuclear localization was performed according to the procedure of (Lin et al., 2018) and statistical analysis was performed using a chi-square test of independence.

### RNA sequencing

#### Isolation of *C. elegans* mRNA

Worms were synchronized, grown and treated with compounds from L4 stage as described. Twenty-four hours later, worms were washed from treatment plates, 3 times in M9 buffer and 2 times in water before being snap frozen in liquid nitrogen. Four samples were grown, each of ∼1000 worms. For isolation of total mRNA, whole worms were homogenized with a 5 mm steel bead using a TissueLyser II (QIAGEN) for 5 min at frequency of 30 times/second. RNA was extracted according to the instructions of the RNaesy Mini Kit (QIAGEN). Contaminating genomic DNA was removed using RNase-Free DNase (QIAGEN). RNA was quantified with a NanoDrop 2000 spectrophotometer (Thermo Scientific; Breda, The Netherlands) and stored at -80°C until use.

#### Library Preparation

RNA libraries were prepared and sequenced with the Illumina platform by Genome Scan (Leiden, The Netherlands). The NEBNext Ultra II Directional RNA Library Prep Kit for Illumina was used to process the sample(s). The sample preparation was performed according to the protocol “NEBNext Ultra II Directional RNA Library Prep Kit for Illumina” (NEB #E7760S/L). Briefly, mRNA was isolated from total RNA using the oligo-dT magnetic beads. After fragmentation of the mRNA, cDNA synthesis was performed. This was used for ligation with the sequencing adapters and PCR amplification of the resulting product. The quality and yield after sample preparation was measured with the Fragment Analyzer. The size of the resulting products was consistent with the expected size distribution (a broad peak between 300-500 bp). Clustering and DNA sequencing using the NovaSeq6000 was performed according to manufacturer’s protocols. A concentration of 1.1 nM of DNA was used. NovaSeq control software NCS v1.6 was used.

#### Read Mapping, Statistical Analyses, and Data Visualization

Reads were subjected to quality control FastQC (Andrews, 2010) trimmed using Trimmomatic v0.32 (Bolger, Lohse, & Usadel, 2014) and aligned to the *C. elegans* genome obtained from Ensembl (wbcel235.v91), using HISAT2 v2.1.0 (Kim, Langmead, & Salzberg, 2015). Counts were obtained using HTSeq (v0.11.0, default parameters) (Anders, Pyl, & Huber, 2015) using the corresponding GTF taking into account the directions of the reads. Statistical analyses were performed using the edgeR v3.26.8 (Robinson, McCarthy, & Smyth, 2010) and limma/voom v 3.40.6 (Ritchie et al., 2015) R packages. All genes with more than 2 counts in at least 3 of the samples were kept. Count data were transformed to log2-counts per million (logCPM), normalized by applying the trimmed mean of M-values method (Robinson et al., 2010) and precision weighted using voom (Law et. al., 2014). Differential expression was assessed using an empirical Bayes moderated t test within limma’s linear model framework including the precision weights estimated by voom (Law et al., 2014; Ritchie et al., 2015). Resulting *p* values were corrected for multiple testing using the Benjamini-Hochberg false discovery rate. Data processing was performed using R v3.6.1 and Bioconductor v3.9. Partial least-squares discriminant analysis (PLS-DA) was performed using mixomics (Rohart et al., 2017) setting a variable of importance (VIP) score of greater than 1 as significant. Resulting *p* values (where applicable) were corrected for multiple testing using the Benjamini-Hochberg false discovery rate. Genes were re-annotated using biomaRt using the Ensembl genome databases (v91). Data visualization was performed using gplots (Warnes et al., 2015) and ggplot2 (Wickham, 2016) selecting colors from RcolorBrewer (Neuwirth, 2014). Functional annotation clustering of differentially expressed genes was performed using the DAVID bioinformatics resource with all default settings (Huang et al., 2009). Clusters with an enrichment score of greater than 1.3 were considered significant. The RNA-seq data are available on GEO under the ID GSE149944.

## Supporting information

SuppTableS1_FOXO3oe_Drug_Hits

SuppTableS3_Atracurium_RNAsequencing

SuppTableS4_Functional_Enrichment_Clusters

## Acknowledgements

The authors thank the *Caenorhabditis* Genetics Center at the University of Minnesota for providing *C. elegans* strains. The CGC is funded by NIH Office of Research Infrastructure Programs (P40 OD010440). We thank Perry D. Moerland for assistance in analysis of the RNA sequencing. Work in the Houtkooper group is financially supported by an ERC Starting grant (no. 638290), a VIDI grant from ZonMw (no. 91715305), and the Velux Stiftung (no. 1063). GEJ is supported by a VENI grant from ZonMw (no. 09150161810014).

## Authors’ Contributions

RLM, RHH and GEJ conceived and designed the project. RLM, SWD and RK performed experiments. RLM, SWD and GEJ analyzed the data. AJ and GEJ performed bioinformatic analyses of the RNA sequencing data. MP, BVS and MSK aided in interpretation of data and provided advice. RLM, GEJ and RHH wrote the manuscript with contributions from all other authors.

## Data Availability Statement

The RNA-sequencing data are available on GEO under the ID GSE149944. All other data are available in the manuscript or supplementary materials. Correspondence and requests for materials should be addressed to the corresponding author GEJ.

---

**Supplemental Table S1: Drug screen hits**

**Supplemental Table S2:** Lifespan Statistics.

| *C. elegans* strain | Treatment | Median lifespan (days) | % Change | Number animals (died/total) | P – value against control group |
| --- | --- | --- | --- | --- | --- |
| **Figure 2 Panel A** |  |  |  |  |  |
| Wild Type (N2)* | DMSO vehicle | 16 |  | 86/100 |  |
| | 50 μM zidovudine | 18 | 31.25% | 76/100 | <0.0001 |
| Wild Type (N2) | DMSO vehicle | 15 |  | 64/100 |  |
| | 50 μM zidovudine | 18 | 20.00% | 69/100 | 0.0024 |
| Wild Type (N2) | DMSO vehicle | 16 |  | 90/100 |  |
| | 50 μM zidovudine | 19 | 18.75% | 80/100 | 0.0001 |
| Wild Type (N2) | DMSO vehicle | 11 |  | 77/100 |  |
| | 50 μM zidovudine | 25 | 127.30% | 76/100 | <0.0001 |
| Wild Type (N2)* | water vehicle | 13 |  | 82/100 |  |
| | 50 μM atracurium | 20 | 58.33% | 81/100 | <0.0001 |
| Wild Type (N2) | water vehicle | 15 |  | 72/100 |  |
| | 50 μM atracurium | 18 | 20.00% | 71/100 | <0.0001 |
| Wild Type (N2) | water vehicle | 13 |  | 85/100 |  |
| | 50 μM atracurium | 16 | 23.08% | 72/100 | 0.0055 |
| Wild Type (N2) | water vehicle | 17 |  | 85/100 |  |
| | 50 μM atracurium | 22 | 22.73% | 76/100 | 0.0002 |
| Wild Type (N2)\$ | water vehicle | 16 | | 77/100 | |
| | 50 μM atracurium | 22 | 37.50% | 63/100 | 0.0004 |
| **Figure 2 Panel B** |  |  |  |  |  |
| *daf-16(mu86)\** | DMSO vehicle | 12 |  | 82/100 |  |
| | 50 μM zidovudine | 19 | 58.33% | 81/100 | <0.0001 |
| *daf-16(mu86)* | DMSO vehicle | 13 |  | 85/100 |  |
| | 50 μM zidovudine | 18 | 38.46% | 69/100 | <0.0001 |
| *daf-16(mu86)* | DMSO vehicle | 10 |  | 84/100 |  |
| | 50 μM zidovudine | 19 | 90.00% | 81/100 | <0.0001 |
| *daf-16(mu86)\** | water vehicle | 13 |  | 82/100 |  |
| | 50 μM atracurium | 13 | 0.00% | 81/100 | 0.0364 |
| *daf-16(mu86)* | water vehicle | 12 |  | 85/100 |  |
| | 50 μM atracurium | 15 | 25.00% | 69/100 | 0.0382 |
| *daf-16(mu86)* | water vehicle | 12 |  | 64/100 |  |
| | 50 μM atracurium | 14 | 7.69% | 81/100 | <0.0001 |
| **Figure 3 Panel G** |  |  |  |  |  |
| Wild Type (N2)* | water vehicle | 17 |  | 85/100 |  |
| | 50 μM atracurium | 22 | 22.73% | 76/100 | 0.0002 |
| Wild Type (N2) | water vehicle | 13 |  | 82/100 |  |
| | 50 μM atracurium | 20 | 58.33% | 81/100 | <0.0001 |
| Wild Type (N2) | water vehicle | 15 |  | 72/100 |  |
| | 50 μM atracurium | 18 | 20.00% | 71/100 | <0.0001 |
| Wild Type (N2) | water vehicle | 13 |  | 85/100 |  |
| | 50 μM atracurium | 16 | 23.08% | 72/100 | 0.0055 |
| Wild Type (N2)<sup>§</sup> | water vehicle<br>50 μM atracurium | 16<br>22 | 37.50% | 77/100<br>63/100 | 0.0004 |
| **Figure 3 Panel H** |  |  |  |  |  |
| *unc-38(e264)\** | water vehicle<br>50 μM atracurium | 15<br>11 | -26.67% | 83/100<br>82/100 | <0.0001 |
| *unc-38(e264)* | water vehicle<br>50 μM atracurium | 13<br>13 | 0.00% | 88/100<br>68/100 | <0.0001 |
| *unc-38(e264)<sup>§</sup>* | water vehicle<br>50 μM atracurium | 16<br>14 | -14.29% | 81/100<br>66/100 | <0.0001 |
| **Figure 3 Panel I** |  |  |  |  |  |
| *unc-38(x20)\** | water vehicle<br>50 μM atracurium | 19<br>19 | 0.00% | 66/100<br>79/100 | 0.1718 |
| *unc-38(x20)* | water vehicle<br>50 μM atracurium | 23<br>21 | -8.70% | 81/100<br>78/100 | 0.0380 |
| *unc-38(x20)<sup>§</sup>* | water vehicle<br>50 μM atracurium | 25<br>25 | 0.00% | 41/100<br>39/100 | 0.0878 |
| **Not shown** |  |  |  |  |  |
| Wild Type (N2) | DMSO vehicle<br>50 μM sildenafil | 15<br>15 | 0.00% | 64/100<br>82/100 | 0.4184 |
| Wild Type (N2) | DMSO vehicle<br>50 μM sildenafil | 16<br>14 | -12.50% | 90/100<br>90/100 | 0.169 |
| Wild Type (N2) | DMSO vehicle<br>50 μM sildenafil | 16<br>13 | -18.75% | 86/100<br>77/100 | 0.0807 |

\*experiment represented in figure
<sup>§</sup>experiment was stopped amid the COVID-19 crisis, though all conditions reached the point of 50% viability

**Supplemental Table S3: RNAseq data**

**Supplemental Table S4:Enrichment cluster analysis**

**Supplemental Table S5:**
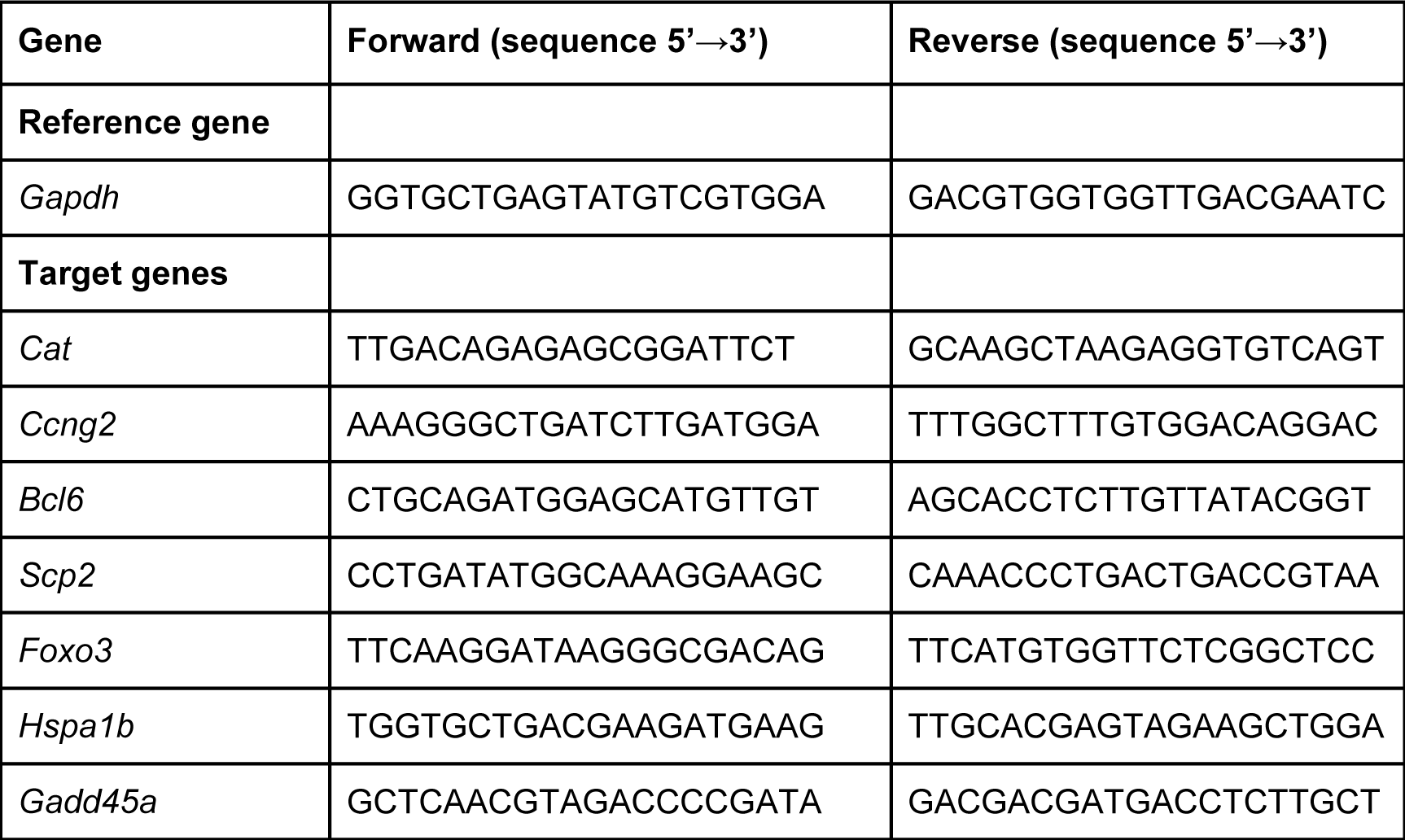
Primers used for HL1 mouse cardiomyocyte qPCR.

